# Image-based Phenotyping and Disease Screening of Multiple Populations for resistance to *Verticillium dahliae* in cultivated strawberry *Fragaria x ananassa*

**DOI:** 10.1101/497107

**Authors:** H.M. Cockerton, B. Li, R. J. Vickerstaff, C. A. Eyre, D. J. Sargent, A. D. Armitage, C. Marina-Montes, A. Garcia, A. J. Passey, D. W. Simpson, R. J. Harrison

**Author notes:** Correspondence: Richard J. Harrison.

## Abstract

*Verticillium dahliae* is a highly detrimental pathogen of soil cultivated strawberry (*Fragaria x ananassa*). Breeding of Verticillium wilt resistance into commercially viable strawberry cultivars can help mitigate the impact of the disease. In this study we describe novel sources of resistance identified in biparental strawberry populations, creating a wealth of data for breeders to exploit. Pathogen-informed experiments have allowed the differentiation of subclade-specific resistance responses, through studying *V. dahliae* subclade II-1 specific resistance in the cultivar ‘Redgauntlet’ and subclade II-2 specific resistance in ‘Fenella’ and ‘Chandler’.

A large-scale low-cost phenotyping platform was developed utilising automated unmanned vehicles and near infrared imaging cameras to assess field-based disease trials. The images were used to calculate disease susceptibility for infected plants through the normalized difference vegetation index score. The automated disease scores showed a strong correlation with the manual scores.

A co-dominant resistant QTL; *FaRVd3D*, present in both ‘Redgauntlet’ and ‘Hapil’ cultivars exhibited a major effect of 18.3 % when the two resistance alleles were combined. Another allele, *FaRVd5D*, identified in the ‘Emily’ cultivar was associated with an increase in Verticillium wilt susceptibility of 17.2%, though whether this allele truly represents a susceptibility factor requires further research, due to the nature of the bi-parental cross.

Markers identified in bi-parental populations were validated across a set of 92 accessions to determine whether they remained closely linked to resistance genes in the wider germplasm. The resistant markers *FaRVd2B* from ‘Redgauntlet’ and *FaRVd6D* from ‘Chandler’ were associated with resistance across the wider germplasm. Furthermore, comparison of imaging versus manual phenotyping revealed the automated platform could identify three out of four disease resistance markers. As such, this automated wilt disease phenotyping platform is considered to be a good, time saving, substitute for manual assessment.

## 3.0 Introduction

*Verticillium dahliae* (Kleb.) is a soilborne plant pathogen which has a large detrimental impact on the yield of soil cultivated strawberry (*Fragaria* x *ananassa*) (Maas, 1998). This ascomycete fungi is particularly problematic due to the longevity of inoculum in the soil whereby the resting propagules, termed microsclerotia, persist for up to 14 years in the absence of a host plant (Schnathorst, 1981). Low inoculum densities of 2 colony forming units per gram of soil can result in complete strawberry crop losses (Harris and Yang, 1996), indicating that strawberry exhibits a very high susceptibility to Verticillium alongside the crops cotton (Paplomatas et al., 1992) and olive when artificially inoculated (Lopez-Escudero & Blanco-Lopez, 2002). Verticillium infects over 200 different dicotyledonous plant species including many horticultural crops and weeds (Bhat and Subbarao, 1999; Woolliams, 1966) meaning that crop rotation is an ineffective form of disease control (Atallah et al., 2011). Effective disease control is also hampered by the absence of curative fungicides and restriction of preventative chemical fumigants due to European regulations (e.g. 91/414/EEC; Colla et al., 2012). Disease resistant germplasm is therefore an essential resource required to combat the pathogen, particularly where countries rely predominantly on soil cultivation systems.

A pathogenesis related protein which catalyzes chitinase from wild tomato has been shown to be effective against *V. dahliae* when transformed into strawberry (Chalavi and Tabaeizadeh, 2003). This mechanism acts before infection therefore indicating very strong resistance as proven by the percentage infection of verticillium in strawberry crowns. Complete resistance has not been observed in natural populations of octoploid strawberry to date. Tolerance, whereby the host is colonised by the fungus but does not exhibit infection symptoms, is frequently observed in strawberry alongside the crop species olive (López-Escudero et al., 2004) potato (Dan et al., 2001), cultivated tomato (Fradin et al., 2009; Chen et al., 2004) and cotton (Bolek et al., 2005; Zhang et al., 2011).

High variation for *V. dahliae* resistance has been observed in Californian strawberry germplasm and empirical selection had led to an increase in resistance (Shaw et al., 1997; Shaw and Gubler, 1996). Studies investigating the general combining ability (GCA) for *V. dahliae* resistance in strawberry, found that four out of ten cultivars had a significant GCA indicating a high transmission of resistance or susceptibility status from parent to progeny. This study suggests that Verticillium wilt resistance is controlled by additive quantitative genetic components (Masny et al., 2014). Furthermore, a significant specific combining ability (SCA) in two crosses indicated that some Verticillium resistance alleles are non-additive (Masny et al., 2014). Previous studies using *in vitro* strawberry have found Verticillium resistance to be controlled by additive genes and in one case a single partially dominant gene (Zebrowska et al., 2006). The study of the bi-parental population ‘Redgauntlet’ x ‘Hapil’ revealed that multiple small effect QTL control *V. dahliae* resistance (Antanaviciute et al., 2015).

Isolate and cultivar specific interactions complicate the description of resistance and must be considered for robust disease resistance breeding. Segregation of *V. dahliae* into six distinct races has been proposed based on the resistance status of different strawberry varieties (Govorova and Govorov, 1997) indicating a complex series of host-pathogen interactions. By contrast, a simpler dissection of *V. dahliae* isolate virulence has been proposed: two subclades of *V. dahliae* have been isolated from UK strawberry; II-1 and II-2, which exhibit different average levels of virulence on the susceptible strawberry cultivar ‘Hapil’ (Fan et al., 2018; Jiménez□Díaz and Olivares□García, 2017).

Single major gene resistance to *V. dahliae* has been identified in tomato, lettuce and cotton; the *Ve 1* host gene, which recognizes the avirulence pathogen effector *VdAve 1*, leads to the separation of *V. dahliae* isolates into two races; those with and without *VdAve 1* (Kawchuk et al., 2001; Hayes et al., 2011; Zhang et al., 2011; de Jonge et al., 2012). Fan et al. (Fan et al., 2018) conclude that there is an absence of the *VdAve 1* gene in *Verticillium dahliae* isolated from UK strawberry. The exclusive infection of strawberry by ‘race 2’ isolates in the UK, despite of the presence of ‘race 1’ isolates in other UK hosts, likely suggests a lack of dispersion of *VdAve1* isolates, rather than selection against *Ave1*, as *VdAve1* isolates were also able to infect strawberry. This reduces the relevance of harnessing *Ve1* mediated resistance in future strawberry breeding.

Platforms for strawberry genotyping have advanced substantially over the last decade (Verma et al., 2017; Bassil et al., 2015), however the low throughput capacity of traditional large scale phenotyping is now the limiting factor restricting pre-breeding research (Mahlein, 2016). Currently, many breeders use manual assessments to quantify the disease resistance status of plants, which is subjective and time consuming. Imaging techniques have been successfully applied to high-throughput plant phenotyping for the past decade (Barbedo, 2013) and with the development of lightweight unmanned aerial vehicles (UAV) for precision agriculture, imaging techniques can be applied to screen large crop areas with centimeter level spatial resolution and accurate positional information (Candiago et al., 2015). Multispectral cameras are lighter than the majority of imaging sensors that can be attached to UAV (Sugiura et al., 2016) they also provide accurate quantification and are a cost effective strategy for disease severity quantification. The most common vegetation index derived from multispectral sensor is the Normalized Difference Vegetation Index (NDVI) where a positive NDVI value indicates healthy green vegetation whilst a negative value indicates the absence of vegetation (Candiago et al., 2015).

In this study, a low-cost UAV with global positioning system and multispectral imaging sensor was implemented as part of a phenotyping platform to measure *Verticillium* wilt resistance in strawberry. We also report a reanalysis of historical data using the ‘Redgauntlet’ and ‘Hapil mapping populations infected with a mixed inoculum of *V. dahliae*, reported by Antanaviciute, (2015) using newly generated SNP data and also test additional progeny of ‘Redgauntlet’ and ‘Hapil against a single isolate from subclade II-1. Furthermore, two additional mapping populations are studied to identify putative resistance loci towards a highly virulent subclade II-2 isolate of *V. dahliae*.

## 4.0 Materials and Methods

### 4.1 Study area and experimental design

Field phenotyping for *V. dahliae* resistance was conducted on three strawberry mapping populations. Mapping populations were produced through crosses between the cultivars ‘Emily’ x ‘Fenella’ (ExF, 181 genotypes), ‘Flamenco’ x ‘Chandler’ (FxC, 140 genotypes) and ‘Redgauntlet’ x ‘Hapil’ (RxH^b^, 160 genotypes). The RxH^b^ cross differs from the population described in previous research, as it is a different set of individuals (Antanaviciute et al., 2015; Sargent et al., 2012). The analysis reported in Antanaviciute et al. (2015) used the original ‘Redgauntlet’ x ‘Hapil’ (RxH^a^) cross and SSR markers. This study integrates the Antanaviciute et al. (2015) phenotypic data where, in contrast to the previous analysis, the Area Under the Disease Progression Curve AUDPC and Best Linear Unbiased Estimate (BLUE) scores were calculated to represent the disease score of each genotype across three years of phenotyping. Use of SNP marker genotyping allowed a more powerful analysis and comparison of resistance markers across populations. The validation experiment utilised 92 accessions selected from across the wider germplasm. Parent and progeny stock plants were maintained in a polytunnel and runners were pinned down into 9 cm pots before planting in ‘Calves Leys’, Aylesford, Kent UK field in autumn 2015 (ExF & FxC) or ‘Rocks Farm’, East Malling, Kent, UK in 2016 (Validation & RxH^b^). Plants were arranged East to West with 64 plants per row at 0.6 m intervals in a randomised block design with 5-10 replicate plants per genotype or accession and parental lines. Black MyPex^®^ was used for weed growth suppression and allowed segregation of plant foliage for image analysis. Plants were rainfed with additional overhead irrigation supplied if required. The pre-existing microsclerotia level was quantified using the Harris method (Harris et al., 1993) and found to be 4.2 cfu g^-1^ soil in ‘Calves Leys’ and 0.9 cfu g^-1^ in ‘Rocks Farm’. To ensure robust disease symptom expression, plants were inoculated with 10 ml of 4 x 10^6^ conidia ml^-1^ into the crown and immediate surrounding soil of each strawberry plant. A single, highly virulent isolate of *V. dahliae* (12008) was used as inoculum in March 2016 for ExF, FxC and March 2017 for germplasm experiments. The isolate, 12008, has been used extensively in work conducted by Soares et al., (2004) and Fan et al., (2018) and represents an isolate from *V. dahliae* subclade II-2, the ‘high virulence subclade’ when inoculated onto strawberry. Plants in the RxH^b^ phenotyping event were inoculated with isolate 12158 from subclade II-1. Weather conditions were 12.2 (±3.7) ^o^C; 76.7 (±8.6) RH% spring 2016, 18.5 (±2.3) ^o^C; 77.4 (±6.3) RH% summer 2016, 13.34 (± 0.49) ^o^C; 74.5 (± 0.76) RH% spring 2017 and 18.36 (±0.62) ^o^C; 75.76 (± 1.12) RH% summer 2017.

### 4.2 Visual assessment of *Verticillium* wilt

Disease scores were recorded five times from June to September at three-week intervals, plants were scored for percentage wilting disease symptoms on a score of 1-9 depending on severity of leaf wilting where a score of 1 denoted a completely healthy plant; 3 denoted 25 % necrotic leaves; 5 denoted 50 % necrotic leaves; 7 denoted 75 % necrotic leaves and 9 denoted 100 % necrosis, a dead plant (Antanaviciute et al., 2015). The AUDPC was calculated across each phenotyping event using the R package “agricolae” (Felipe, 2017) to predict scores for QTL analysis. AUDPC was calculated as below (Shaner and Finney, 1977).

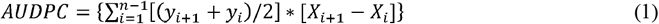

Where y is the disease score, for score i and X represents time in days and n is the number of scoring events. Relative AUDPC (rAUDPC) was calculated by dividing the AUDPC by the number of days after inoculation.

### 4.3 Image acquisition and processing

Arial imaging was conducted, in addition to manual scoring, for 2017 field trials. The 2017 trials were of the RxH^b^ population and the validation set, the experimental field was 45 m x 30 m in size containing approximately 2500 plants (Figure 1a). The UAV platform was a 1.6 kg DJI Flamewheel F450 quadcopter. RGB images were captured using a Canon SX240 HS, 12 MP digital camera. Multi-spectral images with resolution of 1280 x 960 pixels were captured using a MicaSense RedEdge narrow-band multispectral camera (MicaSense, Seattle, Washington). Images were captured at altitude of 30 m at 5 bands including Blue (B: 475 nm center wavelength, 20 nm bandwidth), Green (G: 560 nm, 20 nm), Red (R: 668 nm, 10 nm), Red Edge (RE: 717 nm, 10 nm) and Near Infrared (NIR: 840 nm, 40 nm) were captured simultaneously with the format of 16-bit raw GeoTIFF. Ortho-mosaic images were produced by processing UAV images in Pix4Dmapper Pro software (Pix4D SA, 1015 Lausanne, Switzerland). Two surveys were undertaken of the experimental plot on the 2^nd^ August 2017 at11:00 and the 13^th^ September2017 at12:00. NDVI was calculated as the normalized ratio between near infrared (NIR) and red (R) bands (Potgieter et al., 2017), which is shown in Eq. (2). The diseased:healthy leaf area was calculated based on the green and total plant pixels, which is shown in Eq. (3).

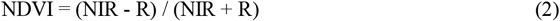

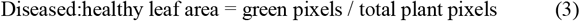

**Figure 1.**
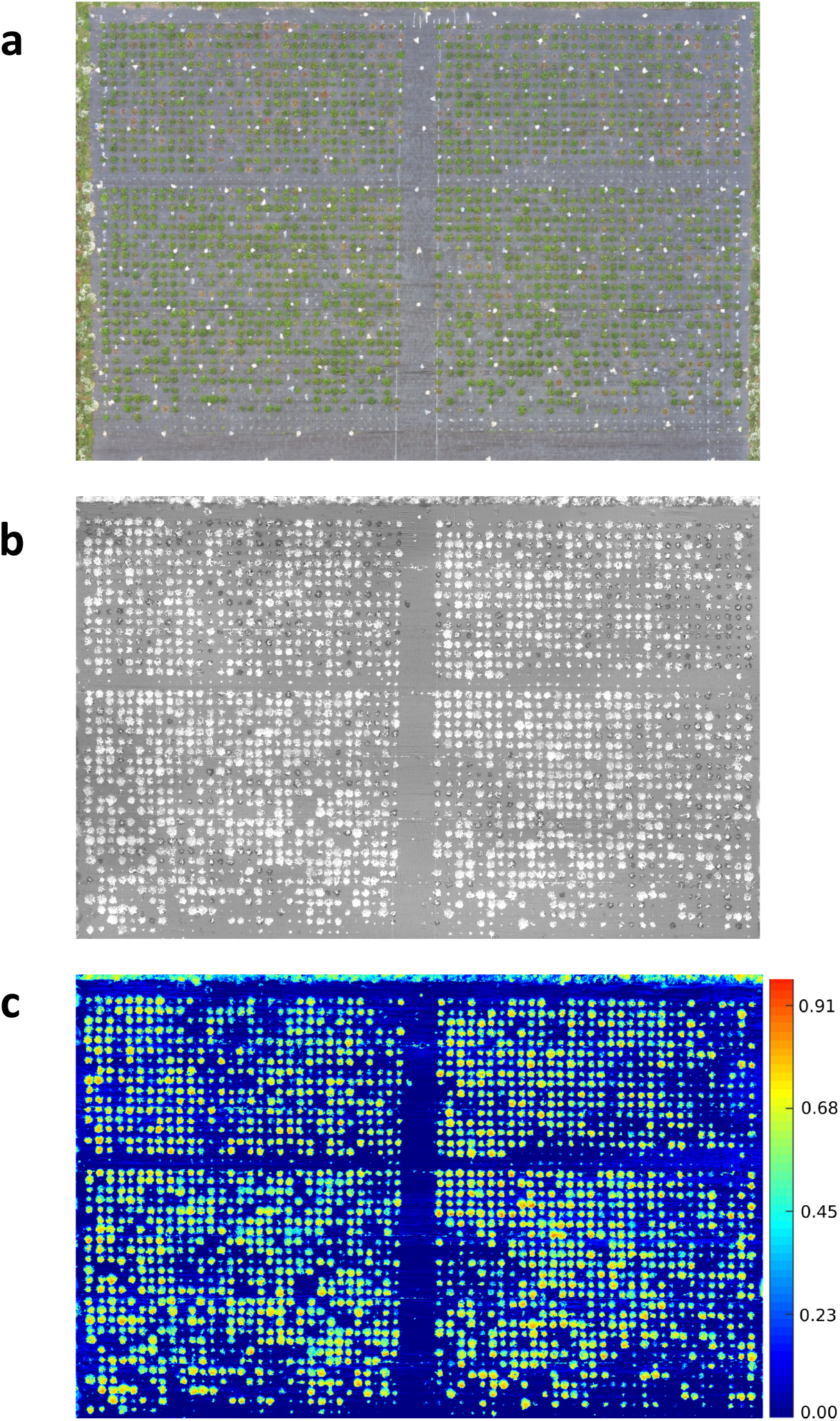
Aerial image taken using UAV of 2017 Verticillium disease field experiments containing the RxH^b^ population and the validation set a) RGB image b) Green: Red ratio mask image of the canopy for each strawberry plant c) Normalised difference vegetation index (NDVI) image with false colour of the validation set and the ‘Redgauntlet’ x ‘Hapil’ (RxH^b^) mapping population.

Bandpass thresholding was applied to obtain the mask image of the whole canopy for each strawberry plant, the green:red band ratio image was found to provide a good contrast between the plant canopy and background (Figure 1b). A semi-automated image analysis software was developed to calculate the average NDVI value for each plant (Figure 1c). Manual selection of a plant on the masked image allows the software to automatically calculate the ratio of total NDVI: total canopy pixel number.

### 4.4 Linkage map generation

The Qiagen DNAeasy plant mini extraction kit (Qiagen Ltd., Manchester, UK) was used to extract DNA from the studied genotypes and accessions according to the manufacturer’s instructions. Biparental populations RxH^a^, ExF and FxC were genotyped using the Affymetrix Istraw90 Axiom^®^array (i90k; Bassil et al., 2015) whereas the population RxH^b^ and validation accessions were genotyped on the streamlined Axiom^®^ IStraw35 384HT array (i35k; Verma et al., 2017). Crosslink was used to generate linkage maps (https://github.com/eastmallingresearch/crosslink) a program developed specifically for polyploid plant species (Vickerstaff and Harrison, 2017). *Fragaria* x *ananassa* chromosome number is denoted by 1-7 and sub-genome number is represented by A-D as specified in van Dijk et al. (2014) and Sargent et al. (2015).

### 4.5 Statistical analysis

For the RxH^a^ historical data the best linear unbiased estimate (BLUE) was calculated using the relative AUDPC for QTL analysis (R package “nlme”, Pinheiro et al., 2017).

For populations phenotyped with both manual and UAV imaging, the Pearson’s correlation coefficient was calculated between the ratio of healthy:diseased leaf area, NDVI and the raw phenotypic disease score at each time point. A combined analysis used the NDVI-AUDPC and healthy:diseased leaf area-AUDPC alongside the AUDPC disease score to determine the efficacy of the drone phenotyping method. Transgressive segregation where progeny wilt phenotype varied more than expected based on parental phenotypes was assessed using a Dunnett’s test.

Disease resistance markers were identified and validated as outlined in Cockerton et al. (2018). Furthermore, inference of whether resistant markers were present across multiple populations and the targeted marker association study was conducted as outlined in Cockerton et al. (2018). Candidate resistance genes were identified in the *Fragaria vesca* genome (assembly v1.1; Shulaev et al., 2011) and screened for the presence of NB-LRR, TM–CC, RLP, RLK (S-type and general) domains and candidate Rosaceous MLO genes (Pessina et al., 2014) following published pipelines (Li et al., 2016). Resistance genes were identified within 100 kb of the significant resistance marker using BEDtools (Quinlan and Hall, 2010). Characterisation of homologous genes in the NCBI database was undertaken through tblastx (Karlin and Altschul, 1993). NB-ARC domains were identified from *F. vesca ab initio* and hybrid gene models using InterProScan (Quevillon et al., 2005). Significant association of NBS and NB-ARC domains with focal markers was tested through assessing their occurrence within 100 kb of 25 randomly sampled markers from across the four populations over 10,000 permutations.

## 5.0 Results

### 5.1 Resistance to isolates varies between cultivars

The cultivar ‘Redgauntlet’ exhibits tolerance to the subclade II-1 isolate, 12158 and moderate tolerance to the subclade II-2 isolate 12008 however the cultivar ‘Hapil’ is susceptible to the isolates from both subclades (Sup Figure 1). The cultivars ‘Fenella’, ‘Flamenco’ and ‘Chandler’ are highly tolerant to the *V. dahliae* subclade II-2 isolate whereas ‘Emily’ is highly susceptible (Sup Figure 2).

### 5.2 The ‘Flamenco’ x ‘Chandler’ linkage map

The newly generated FxC linkage map (Sup table 1) has an average genetic distance between markers of 0.3 cM which is a lower average gap than ExF and RxH^a^ (Cockerton et al. 2018), however there are 10 gaps greater than 20 cM and linkage groups 2C, 3C, 6C and 6D each resolved into two linkage groups. FxC linkage information was used as one of the five populations to construct the consensus map. All reported marker positions listed in this study are based on the position in the consensus map.

### 5.4 Comparison of automated and manual phenotyping methods

The proportion of diseased: healthy leaf area and the NDVI values were assessed at discrete time points and over time (Figure 1). A strong negative correlation was observed between the manual disease scores and diseased: healthy leaf area (Figure 2), with a stronger relationship observed between the manual disease score and NDVI for validation and RxH^b^ phenotyping events 2/8/17 (*r* = 0.78, *p* < 0.001) and 13/9/17 (*r*= 0.78, *p* < 0.001) and also between the manual AUDPC and NDVI-AUDPC (*r* = 0.85, *p* > 0.001; Figure 2).

**Figure 2.**
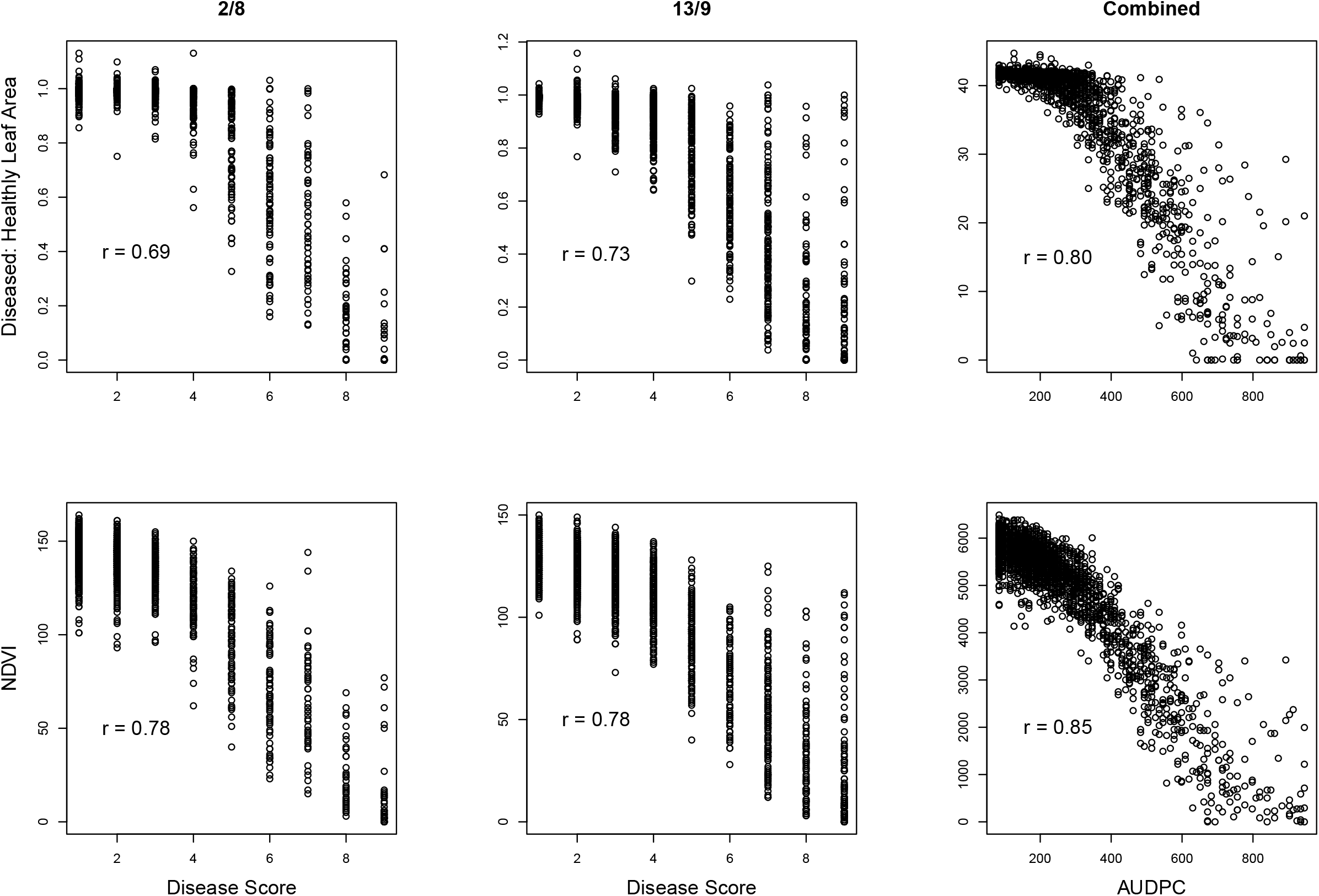
Correlation of disease scores measured through manual and automated techniques. Manual disease score against Diseased: Healthy and Normalised difference vegetation index (NDVI) leaf area for observation time points on 2^nd^ August (2/8) and 13^th^ September (13/9). Area under the disease progression curve calculated for combine Diseased: Healthy score and NDVI. Strawberry plants inoculated with *Verticillium dahliae* across the validation set the ‘Redgauntlet’ x ‘Hapil’ (RxH^b^) mapping population. *r* = pearson’s correlation values.

Significance values for focal SNPs predicted by either automated and manual phenotyping follow the same patterns across the strawberry genome (Figure 3) and three out of four resistance markers were successfully identified in the automated phenotyping QTL analysis (Figure 4).

**Figure 3.**
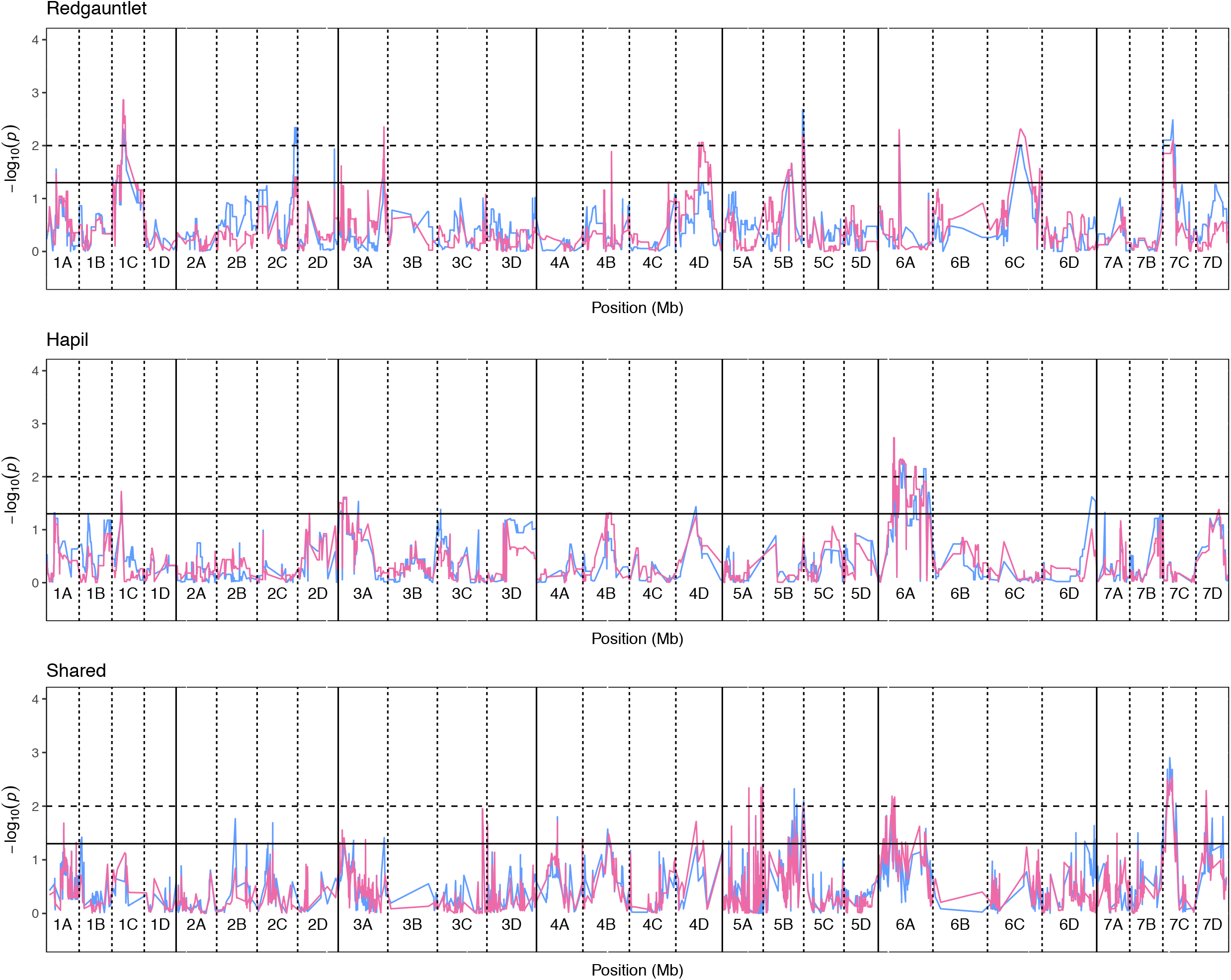
Kruskal-Wallis −log_10_ p-values denoting the association of single nucleotide polymorphisms with strawberry *Verticillium dahliae* automated and manual disease scores at each position in the octoploid strawberry genome in cM. Panels represent markers segregating in ‘Redgauntlet’, ‘Hapil’ and both parents. Labels 1A-7D denote the 28 linkage groups. Solid horizontal line is *p= 0.05*, dashed horizontal line is *p= 0.01*. Colour denotes phenotype measure blue-manual AUDPC, pink- NDVI-AUDPC.

**Figure 4.**
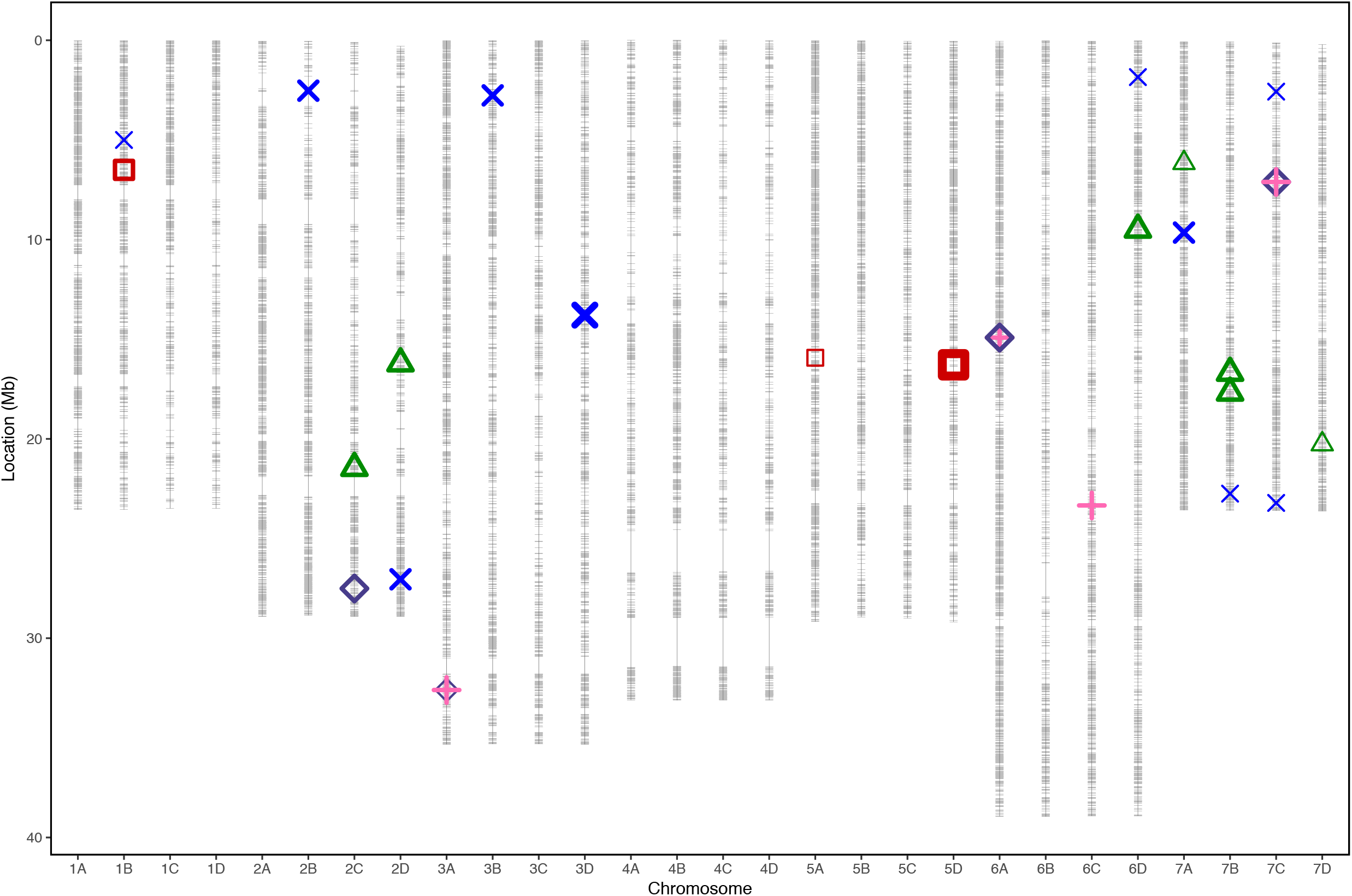
Linkage map displaying 35154 marker positions (grey) in Mb for 28 linkage groups of octoploid strawberry (1A-7D) marker positions scaled to the *F. vesca* genome. Quantitative trait loci associated with strawberry *Verticillium dahliae* disease resistance locations from each phenotyping event represented by points for ‘Emily’ x ‘Fenella’ (red squares), ‘Redgauntlet’ x ‘Hapil’ RxH^a^ (blue cross), RxH^b^ manual scores (pink plus) and RxH^b^ automatic scores (purple diamond) and ‘Flamenco’ x ‘Chandler (green triangle). Points are weighted based on significance with thicker lines representing greater significance.

### 5.3 QTL mapping in four bi-parental populations

In total, four populations were assessed for resistance to *V. dahliae*. Twenty-five focal markers for *V. dahliae* resistance were identified in the RxH^a^, RxH^b^, ExF and FxC populations of strawberry (Figure 3–7, Table 1). Twelve of these focal markers were considered to have a moderate effect with greater than 10 percent impact on disease score across the population. When comparing the observed versus expected disease scores the coefficients of determination, the focal markers explain between 25% and 68% of the observed mean disease scores between progeny members (Table 1).

**Figure 5.**
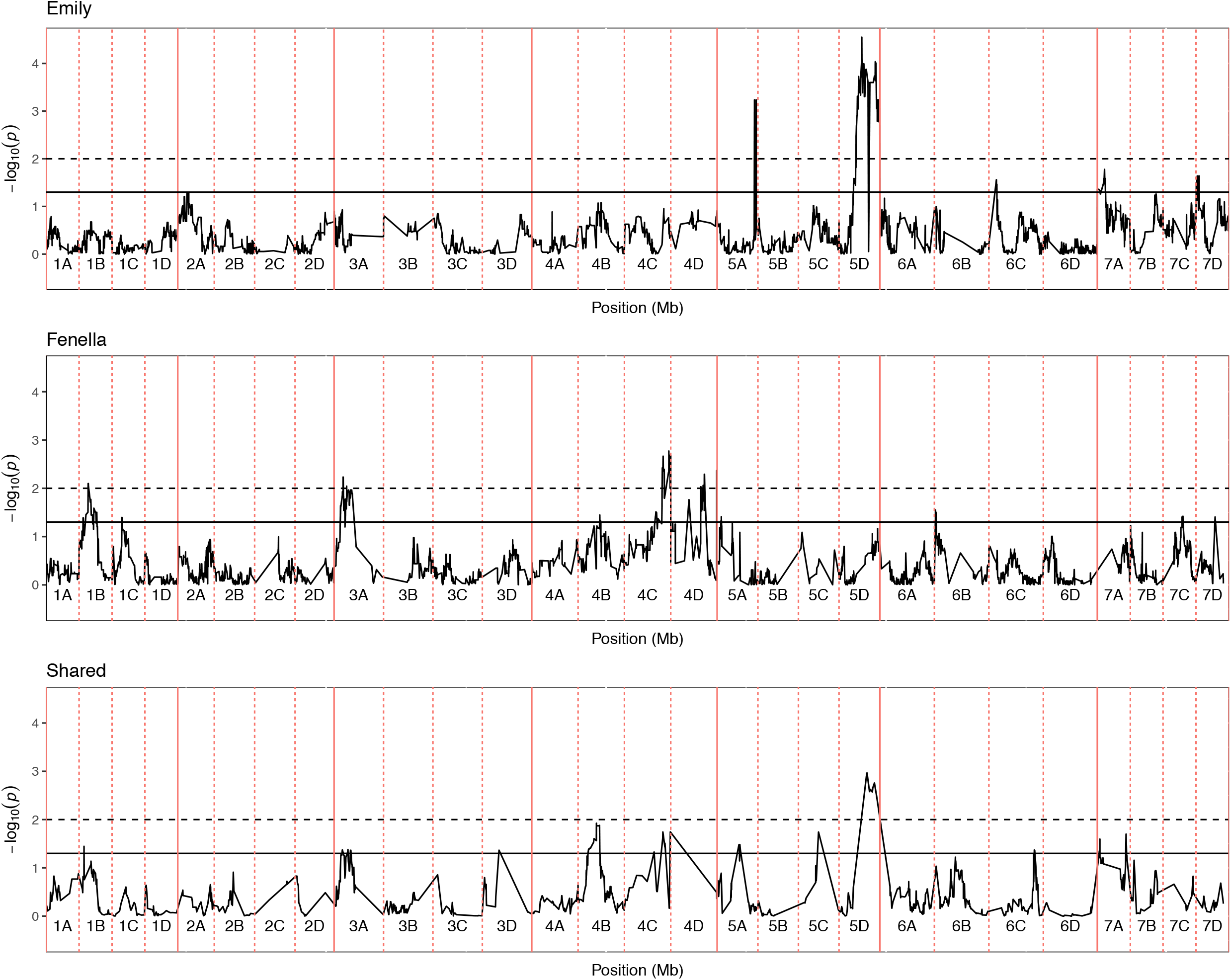
Kruskal-Wallis −log_10_ p-values denoting the association of single nucleotide polymorphism with strawberry *Verticillium dahliae* disease scores at each position in the octoploid strawberry genome in cM. Panels represent markers segregating in ‘Emily’, ‘Fenella’ and both parents. Labels 1A-7D denote the 28 linkage groups. Solid horizontal line is *p= 0.05*, dashed horizontal line is *p= 0.01*.

**Figure 6.**
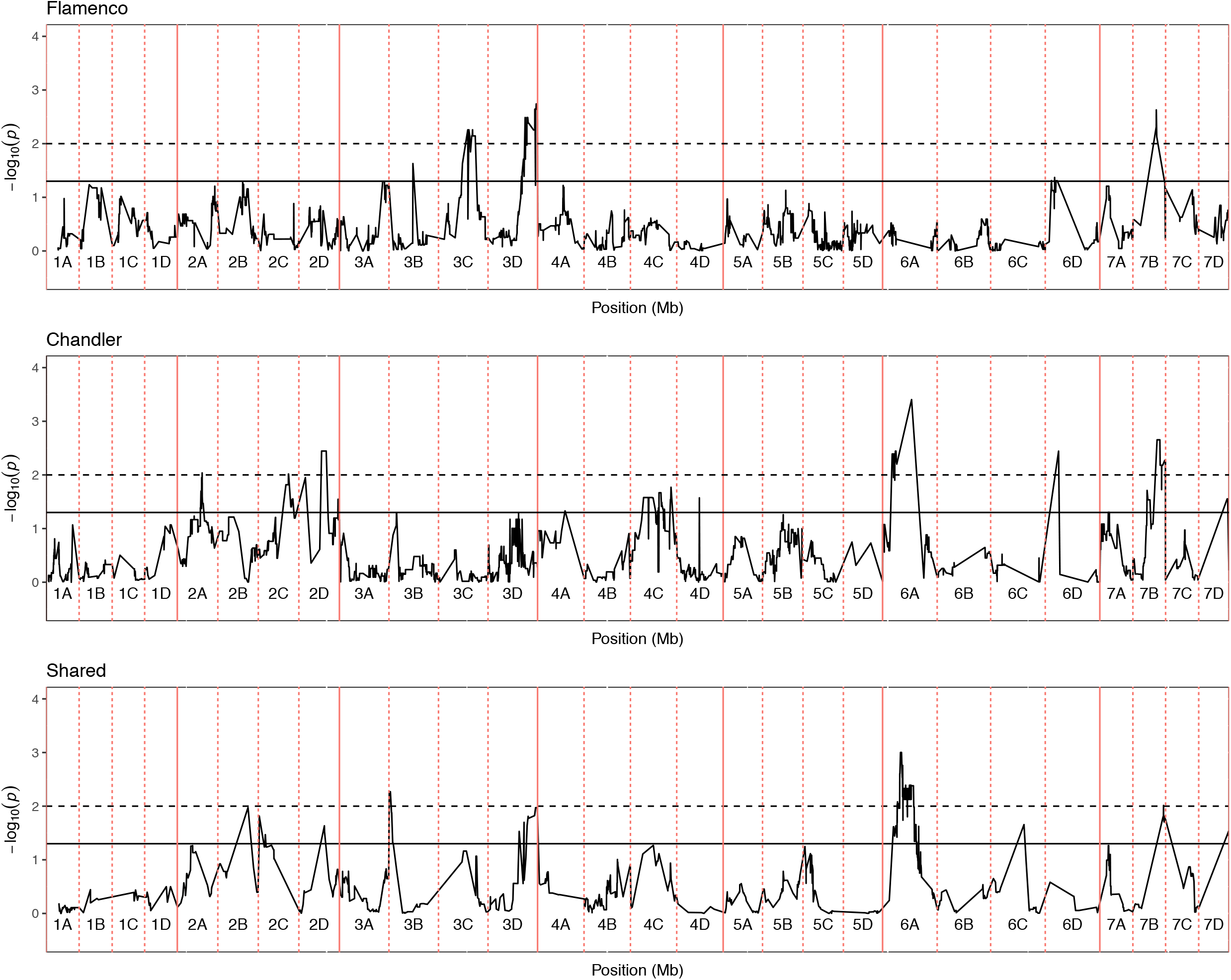
Kruskal-Wallis −log_10_ p-values denoting the association of single nucleotide polymorphism with strawberry *Verticillium dahliae* disease scores at each position in the octoploid strawberry genome in cM. Panels represent markers segregating in ‘Flamenco’, ‘Chandler’ and both parents. Labels 1A-7D denote the 28 linkage groups. Solid horizontal line is *p= 0.05*, dashed horizontal line is *p= 0.01*.

**Figure 7.**
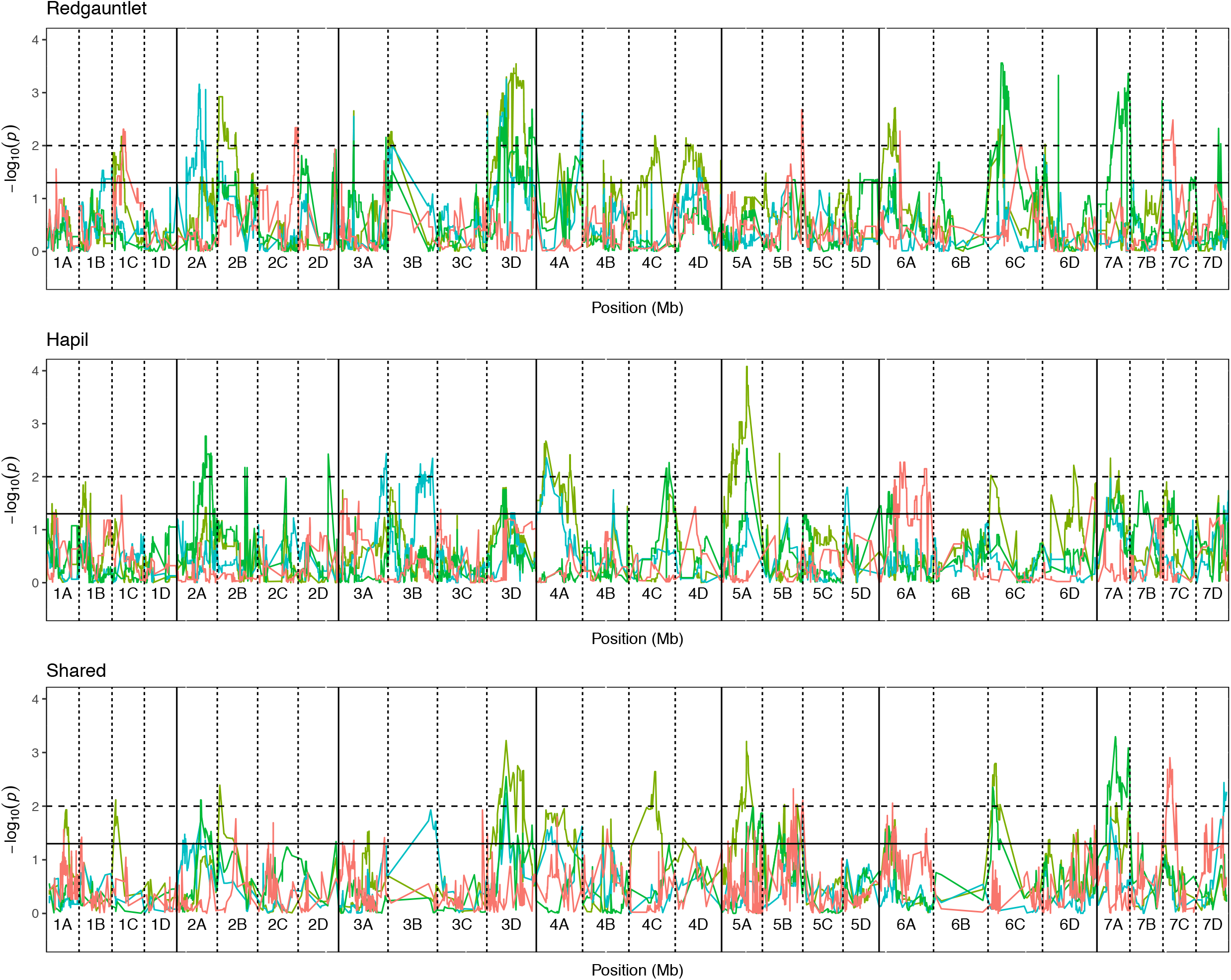
Kruskal-Wallis −log_10_ p-values denoting the association of single nucleotide polymorphism with strawberry *Verticillium dahliae* disease scores at each position in the octoploid strawberry genome in cM. Panels represent markers segregating in ‘Redgauntlet’, ‘Hapil’ and both parents. Labels 1A-7D denote the 28 linkage groups. Solid horizontal line is *p= 0.05*, dashed horizontal line is *p= 0.01*. Colour denotes phenotyping event blue- 2009, lime- 2010, green- 2011, orange- 2017.

**Table 1.** Model parameters for the predictive linear model for each phenotyping event. Predicted versus observed disease scores or each genotype within the population. R^2^ is the coefficient of determination, df are the degrees of freedom associated with the F statistic, the numerator is associated with model parameter number. H^2^ is the broad sense heritability. (A) denotes the automated disease assessment.

| Mapping population | Year | Isolate | R<sup>2</sup> | df | F value | p value | RSE | H<sup>2</sup> |
| --- | --- | --- | --- | --- | --- | --- | --- | --- |
| ExF | 2016 | 12008 | 0.25 | 2,129 | 14.17 | 4.8 x 10<sup>-8</sup> | 69.48 | 0.08 |
| FxC | 2016 | 12008 | 0.52 | 7,71 | 10.97 | 2.6 x 10<sup>-9</sup> | 63.50 | 0.18 |
| RxH<sup>a</sup> | 2009 | Mixed | 0.69 | 15,110 | 16.25 | 2.2 x 10<sup>-16</sup> | 103.80 | 0.45 |
| RxH<sup>a</sup> | 2010 | Mixed | 0.46 | 9,111 | 51.61 | 1.4 x 10<sup>-11</sup> | 51.61 | 0.17 |
| RxH<sup>a</sup> | 2011 | Mixed | 0.48 | 8,118 | 13.64 | 7.4 x 10<sup>-14</sup> | 86.56 | 0.27 |
| RxH<sup>a</sup> | BLUE | Mixed | 0.53 | 11,127 | 13.1 | 2.3 x 10<sup>-16</sup> | 79.13 | NA |
| RxH<sup>b</sup> | 2017 | 12158 | 0.43 | 4,58 | 11.15 | 8.9 x 10<sup>-7</sup> | 50.70 | 0.10 |
| RxH<sup>b</sup> (A) | 2017 | 12158 | 0.42 | 4,58 | 10.58 | 1.7 x 10<sup>-6</sup> | 389.80 | 0.12 |

### 5.5 Resistance markers found close to neighbouring resistance genes

A total of 14 out of 25 markers identified were located within 100 kb of a putative resistance gene found in *F. vesca* (Table 2), each of which indicating a potential target for further study. Twelve resistance markers were found to be within 100 kb of a putative resistance gene containing a nucleotide binding site (NBS). NBS containing genes were more frequently associated with resistance focal SNPs than markers selected at random (*p*= 0.0073 – 0.0017, *n*=10,000; Sup Figure 3). Nine of the twelve NBS genes contained an NB-ARC domain, however NB-ARC domain containing genes were not more frequently associated with resistance focal SNPs than markers selected at random (*p*= 0.066 − 0.024, *n*=10,000; Sup Figure 4).

**Table 2.** Focal single nucleotide polymorphisms linked with each quantitative trait loci associated with strawberry *Verticillium dahliae* disease resistance identified through the Kruskal-Wallis analysis. Closest resistance gene reported within 100 kb if applicable.

| QTL Name | Linkage Group | Closest SNP_i90k | Position (Mb) | p value | k | Sig | Parent | Percentage change | Closest R/S gene (bp) | Type of gene | Gene name | Number R/S genes 100 kb | Population |
| --- | --- | --- | --- | --- | --- | --- | --- | --- | --- | --- | --- | --- | --- |
| *FaRVd5D* | 5D | Affx-88867578 | 16.3 | 2.8 x 10<sup>-5</sup> | 17.5 | **** | Emily | 17.2 | NA | NA | NA | 0 | ExF |
| *FaRVd1B* | 1B | Affx-88814091 | 6.5 | 8.0 x 10<sup>-3</sup> | 7.0 | ** | Fenella | 10.9 | NA | NA | NA | 0 | ExF |
| *FaRVd5A* | 5A | Affx-88865854 | 1.6 | 3.3 x 10<sup>-2</sup> | 8.7 | * | Emily and Fenella | -10.1 | NA | NA | NA | 0 | ExF |
| *FaRVd7B* | 7B | Affx-88898083 | 16.7 | 2.4 x 10<sup>-3</sup> | 9.3 | ** | Flamenco | 9.3 | 9853 | NBS | maker-LG7-exonerate_protein2genome-gene-152.135-mRNA-1 | 6 | FxC |
| *FaRVd2C* | 2C | Affx-88826960 | 21.5 | 3.3 x 10<sup>-3</sup> | 6.7 | ** | Chandler | 10.9 | 34428 | NBS | maker-LG2-augustus-gene-170.171-mRNA-1 | 3 | FxC |
| *FaRVd2D* | 2D | Affx-88821311 | 16.2 | 3.3 x 10<sup>-3</sup> | 8.5 | ** | Chandler | 9.5 | NA | NA | NA | 0 | FxC |
| *FaRVd7B2* | 7B | Affx-88898195 | 17.7 | 3.3 x 10<sup>-3</sup> | 9.4 | ** | Chandler | -9.8 | 25189 | RLK | mrna26334.1-v1.0-hybrid | 1 | FxC |
| *FaRVd7A* | 7A | Affx-88894946 | 6.2 | 3.3 x 10<sup>-2</sup> | 3.8 | * | Chandler | 9.4 | 16557 | NBS | maker-LG7-exonerate_protein2genome-gene-88.76-mRNA-1 | 2 | FxC |
| *FaRVd6D* | 6D | Affx-88877237 | 9.5 | 3.3 x 10<sup>-3</sup> | 8.5 | ** | Chandler | -13.2 | 9934 | NBS | mrna09586.1-v1.0-hybrid | 3 | FxC |
| *FaRVd7D* | 7D | Affx-88900355 | 20.3 | 3.3 x 10<sup>-2</sup> | 4.8 | * | Chandler | 10.7 | 8101 | NBS | mrna34123.1-v1.0-hybrid | 1 | FxC |
| *FaRVd6A* | 6A | Affx-88902601 | 14.9 | 3.3 x 10<sup>-3</sup> | 7.8 | ** | Redgauntlet | 15.2 (-5.0) | NA | NA | NA | 0 | RxH<sup>b</sup><br>Manual & Automated |
| *FaRVd2C2* | 2C | Affx-88828094 | 27.5 | 4.6 x 10<sup>-3</sup> | 8.0 | ** | Redgauntlet | -15.5 | 13390 | NBS | genemark-LG2-processed-gene-188.73-mRNA-1 | 2 | RxH<sup>b</sup><br>Manual |
| *FaRVd3A* | 3A | Affx-88845095 | 32.6 | 1.9 x 10<sup>-2</sup> | 5.5 | * | Redgauntlet | 13.1 (-5.0) | NA | NA | NA | 0 | RxH<sup>b</sup><br>Manual & Automated |
| *FaRVd7C* | 7C | Affx-88895402 | 7.1 | 3.3 x 10<sup>-3</sup> | 8.6 | ** | Redgauntlet | -14.7 (5.9) | 40246 | NBS | maker-LG7-augustus-gene-98.128-mRNA-1 | 1 | RxH<sup>b</sup><br>Manual & Automated |
| *FaRVd6C* | 6C | Affx-88883592 | 23.3 | 4.9 x 10<sup>-3</sup> | 7.9 | ** | Redgauntlet | (-5.3) | NA | NA | NA | 0 | RxH<sup>b</sup><br>Automated |
| *FaRVd6D2* | 6D | Affx-88818910 | 1.8 | 1.1 x 10<sup>-2</sup> | 6.5 | * | Redgauntlet | 6.0 | 13492* | NBS* | maker-LG1-snap-gene-197.206-mRNA-1* | 2* | RxH<sup>a</sup> |
| *FaRVd2B* | 2B | Affx-88822931 | 2.5 | $2.7 \times 10^{-3}$ | 9.0 | ** | Redgauntlet | 6.6 | NA | NA | NA | 0 | RxH<sup>a</sup> |
| *FaRVd2D2* | 2D | Affx-88828415 | 27.0 | $4.2 \times 10^{-3}$ | 8.2 | ** | Redgauntlet | -10.0 | NA | NA | NA | 0 | RxH<sup>a</sup> |
| *FaRVd3D* | 3D | Affx-88836863 | 13.8 | $8.8 \times 10^{-4}$ | 16.5 | *** | Redgauntlet and Hapil | -18.3- both alleles | 82711 | MLO-homolog | mrna31264.1-v1.0-hybrid | 16 (S) | RxH<sup>a</sup> |
| *FaRVd3B* | 3B | Affx-88833107 | 2.8 | $7.9 \times 10^{-3}$ | 7.1 | ** | Redgauntlet | 8.4 | 31622 | NBS | genemark-LG3-processed-gene-23.50-mRNA-1 | 2 | RxH<sup>a</sup> |
| *FaRVd7B3* | 7B | Affx-88900983 | 22.7 | $1.8 \times 10^{-2}$ | 5.6 | * | Redgauntlet | 7.2 | 26652 | NBS | maker-LG7-augustus-gene-201.144-mRNA-1 | 4 | RxH<sup>a</sup> |
| *FaRVd7C2* | 7C | Affx-88900732 | 23.2 | $3.6 \times 10^{-2}$ | 4.4 | * | Redgauntlet | 6.2 | 17146 | NBS | mrna12407.1-v1.0-hybrid | 2 | RxH<sup>a</sup> |
| *FaRVd1B2* | 1B | Affx-88813017 | 5.0 | $3.4 \times 10^{-2}$ | 4.5 | * | Hapil | -9.4 | NA | NA | NA | 0 | RxH<sup>a</sup> |
| *FaRVd7C4* | 7C | Affx-88894332 | 2.6 | $3.4 \times 10^{-2}$ | 4.5 | * | Hapil | -7.7 | NA | NA | NA | 0 | RxH<sup>a</sup> |
| *FaRVd7A2* | 7A | Affx-88895680 | 9.7 | $4.2 \times 10^{-3}$ | 8.2 | ** | Hapil | 5.8 | 52659 | NBS | mrna04795.1-v1.0-hybrid | 2 | RxH<sup>a</sup> |

### 5.6 Improved QTL identification with SNP data

Newly generated SNP data has allowed the analysis of the RxH^a^ mapping population wilt phenotypic data reported in Antanaviciute, (2015). Our new analyses identified four alleles on the same linkage groups as those previously reported (Antanaviciute et al., 2015) and six novel resistance QTL. The original SSR markers associated with wilt resistance all mapped to the same chromosome number as originally reported, however different sub-genomes were assigned when following the linkage group nomenclature stipulated by van Dijk et al. (2014) and Sargent et al. (2015). The QTL *RVd1* maps to linkage group 3B and is located 2.7 Mb from the *FaRVd3B* SNP marker. *RVd3* maps to linkage group 7A and is 6.4 Mb from *FaVd7A2. RVd7* maps to linkage group 2D and is 1.2 Mb away from *FaRVd2D2. RVd4-M1* mapped to linkage group 2B however it is not considered to represent the same QTL as *FaRVd2B* as it was mapped 12.6 Mb away. The RxH^a^ SNP strawberry map has a greater density of segregating loci (3451) than the SSR map (1133) therefore the SNP data allows greater accuracy of QTL mapping which, when combined with the consensus map, assists the comparison of alleles to other phenotyped populations. Discrepancies between the two analyses can be explained by the removal of 13 rogue individuals, the use of AUDPC phenotyping measure and BLUE calculated across multiple years of phenotyping, in comparison the original analysis used data from the single most heritable scoring event for each year.

### 5.7 Overlap of resistance markers between cultivars

The resistance marker *FaRVd7B* identified in ‘Flamenco’ and *FaRVd7B2* in ‘Chandler’ are 9.5 cM and 1 Mb apart however the analysis of haploblocks revealed that these two markers represent discrete resistance loci present on different haplotypes (Figure 8). Shared markers indicate that the resistance marker from ‘Flamenco’ and ‘Chandler’ contribute to resistance in an additive fashion. The haploblock representing the marker *FaRVd7B* is associated with resistance in ‘Flamenco’ is also present in ‘Chandler’ however, it is associated with susceptibility. This low transferability indicates that marker tagging the resistance haplotype *FaRVd7B* is not in linkage disequilibrium with the resistance gene.

**Figure 8.**
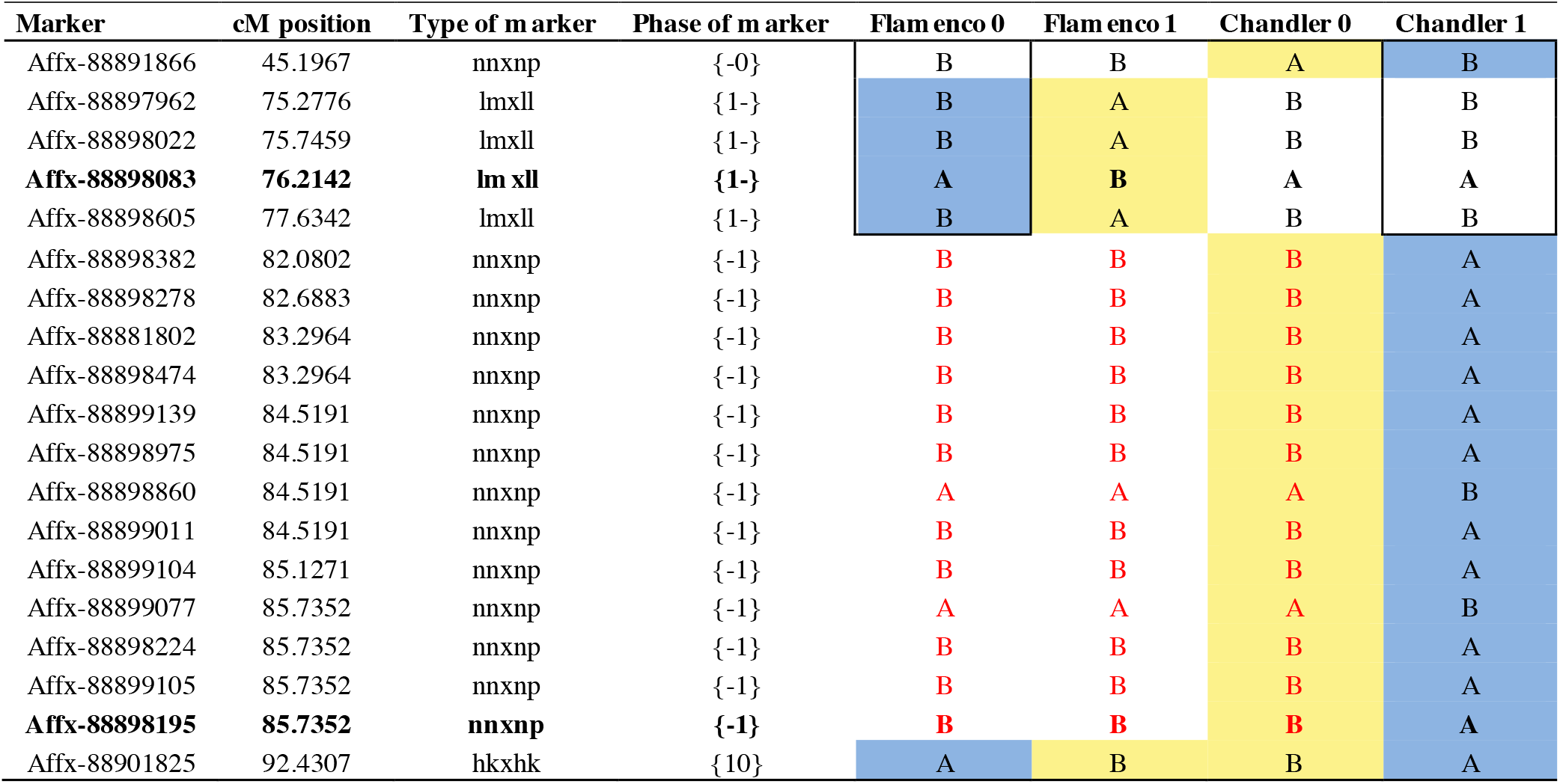
Parental marker on linkage group 7B for phase 0 and 1. Bold text represents focal SNP markers. Red text represents a shared haploblock. A blue background denotes markers associated with resistance and yellow background denotes markers associated with susceptibility.

The resistance markers *FaRVd1B* and *FaRVd1B2* from population RxH^a^ and ExF, respectively, are positioned 1.5 Mb apart on linkage group 1B (Figure 4). Comparison of haploblocks across the two-populations allowed us to determine whether the two markers represent the same resistance allele. Although the focal markers are reciprocally monomorphic, analysis of shared polymorphic neighboring markers indicated that the resistant markers are present on different haplotypes and therefore represent discrete resistance loci.

No overlap in markers was observed between resistance loci identified between the RxH^a^ combined analysis and the RxH^b^ populations screened with mixed inoculum and subclade II-2 inoculum, respectively. By contrast, the marker Affx-88837276 identified on linkage group 7C was 0.72 Mb from the focal marker identified in the RxH^a^ 2011 phenotyping event.

The aforementioned co-dominant shared marker, *FaRVd3D*, was identified in both ‘Redgauntlet’ and ‘Hapil’ cultivars; this is a shared resistance QTL between the two cultivars. The shared marker *FaRVd5A* identified a resistance allele present in both ‘Emily’ and ‘Fenella’ cultivars whereby homozygous genotypes containing two resistance alleles are required to observe significant levels of resistance.

### 5.8 Validation of two resistance markers across the wider germplasm

Identification of substitute i35k SNPs co-localising with focal SNPs identified in the i90k biparental analysis allowed focal markers to be screened across the wider germplasm (Sup Figure 5). Two of the focal SNPs identified in the bi-parental studies maintained a strong association with resistance across the validation accessions. *FaRVd2B* identified in ‘Redgauntlet’ (*X^2^*_(4,5;1)_ = 5.72; *p* = 0.017) and *FaRVd6D* identified in ‘Chandler’ (*X^2^*_(4,6;2)_ = 7.47; *p* = 0.024) explained 13.4 % and 2.2% of the variation in disease scores observed in the validation germplasm, respectively.

## Discussion

### Description of resistance

Multiple sources of resistance to Verticillium wilt were observed across six strawberry cultivars indicating a wealth of genetic resources that can be exploited by breeders. Similar studies have also found multiple sources of resistance to *V. dahliae* in strawberry, and all alleles were found to be dominant (Govorova and Govorov, 1997). We observe that the dominantly inherited allele *FaRV5D* is associated with an increase in susceptibility (Figure 9). It is impossible to know from this work whether *FaRV5D* is a susceptibility factor or whether it represents an additive resistance allele which is in repulsion to the identified marker. Further work, through selfing ‘Emily’, or through crossing heterozygous ExF progeny would generate the missing homozygous class and reveal the inheritance of this resistance allele more clearly. Should *FaRV5D* be found to represent a recessive resistance allele it could prove a valuable tool for strawberry breeders. Resistant homologues of susceptibility factors have been shown to be highly robust, with exploitation lasting for 50 years in the field (Pavan et al. 2010; Kang et al. 2005), therefore utilization of such a resistance incidence could prove a highly robust strategy to prevent against Verticillium infection.

**Figure 9.**
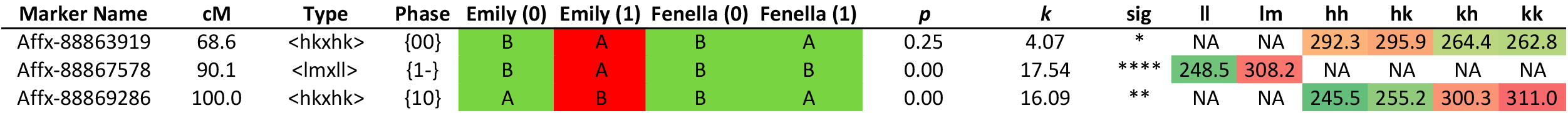
Phasing and marker effect sizes for the *FaRVd5D* focal SNP and neighbouring shared markers. Parental phased haploblocks for linkage group 5D represented in “Emily 0”, “Emily 1”, “Fenella 0” and “Fenella 1” columns, Red haplotype associated with susceptibility. Grand means for each marker class represented under marker classes denoted “ll”, “lm”, “hh”,”hk”, “kh” and “kk”. cM – centimorgan distance along linkage group 5D, *p* – probability for the *k*- Kruskal-Wallace test statistic testing differences between marker classes.

A resistance QTL was identified in both ‘Redgauntlet’ and ‘Hapil’ cultivars on linkage group 3D. Analysis of parental and shared markers in this region indicated resistance alleles from both parents co-localised to the same location and thus represent the same QTL. This QTL was termed *FaRVd3D* and could be best represented by haploblocks phase 1 ‘Hapil’ and phase 0 ‘Redgauntlet’. Both alleles are required in order to observe the greatest combined resistance effect thus indicating that this QTL was inherited in a co-dominant fashion.

Genotypes exhibit large variation in the disease response when compared to variation across genotypes. The variation is represented by large standard error values (Sup. Figure 2) and the corresponding low broad sense heritability values (Table 1). The high correlation between automated and manual phenotyping values, validates the manual phenotypic scores. We can therefore conclude that the large variation associated with disease score reflects the truly variable nature of the verticillium disease responses in strawberry. This within-genotype variation has been observed previously and as such, high replication of genotypes in verticillium trials (*n*=10) mitigates this large variation and results in a greater phenotyping accuracy.

### Transgressive segregation

Transgressive segregation towards susceptibility was observed in the FxC population (Sup Figure 2) with 7.5% of progeny exhibiting a significantly higher disease symptoms than that of the parents. The parental cultivars ‘Flamenco’ and ‘Chandler’ are related, namely ‘Chandler’ is the grandparent of ‘Flamenco’. Reports that inbreeding results in increased susceptibility to plant diseases (Watt et al., 2013) alongside negative implications on other traits (Maas and Galleta, 1996) support the observation of increased susceptibility after crossing two related parents. Transgressive segregation towards susceptibility indicates that the two parental lines contain different resistance alleles (Geiger and Heun, 1989), indeed we do not identify any shared markers or loci between the two parents, however only one significant marker was identified in ‘Flamenco’. Nonetheless, this cross allowed the identification of a number of focal SNPs for further investigation.

### Limitations of the i35k phenotyping platform

Phenotypic data was re-analysed using the subset of i90k markers represented on the i35 SNP chip. The subset of i35k markers were associated with a slightly reduced power to detect resistance markers (Sup Figure 5). A surrogate i35k SNP marker could not be elucidated for *FaRVd6D.* The i35k focal SNP representing the *FaRVd5D* shifted 9.2 Mb and *FaRVd1B2* had shifted 2.0 Mb. However, the remaining focal markers were detected within 0.8 Mb or less of the i90k focal SNP. The validation set of 92 cultivars was phenotyped using the streamlined i35k SNP chip. A targeted marker association analysis using the validation phenotyping event did not pull out any resistance markers, typically genome wide analysis requires a greater genotype number. Either a greater density of markers or a greater number of genotypes may allow the identification of resistance QTL present across the wider germplasm.

### Environmental factors

Variation in weather conditions can lead to variation in disease severity between years (Talboys and Bennett, 1969), the variation in disease development may explain differences observed in RxH^a^ phenotyping events (Keyworth and Bennett, 1951). Where variation in disease susceptibility was observed in cultivars of ‘Earliglow’, ‘Howard 17’ and ‘Bounty’ across different publications (Vining et al., 2015) this may be due to environmental variation or variation in the isolates subclade used for inoculations.

### NBS genes may contribute to Verticillium resistance

A high proportion of the resistance focal SNPs were associated with nucleotide binding site (NBS) resistance genes indicating that NB-LRR mediated signalling may play a role in strawberry Verticillium resistance. NBS genes have been implicated in Verticillium resistance in other host systems. Seven TIR-NBS-LRR resistance genes were observed to be up-regulated in *Arabidopsis thaliana* 24 hours after verticillium co-culturing (Scholz et al. 2018) again indicating NBS-LRRs may play a role in Verticillium resistance. The NBS resistance gene *GbaNA1* was found to control disease resistance to Verticillium in cotton and also confer resistance when transformed into *A. thaliana* (Li et al. 2018a; Li et al. 2018b). A positive correlation between the number of Verticillium and Fusarium wilt resistance QTL and NBS genes was observed on subgenome A of Cotton (Zhang et al. 2015). The most convincing evidence for the existence of Verticillium specific nuclear interactions can be observed through the pathogen effector *VdSCP7* which was found to localise at the host nucleus and modulate effector triggered immunity in cotton (Zhang et al. 2017). Of the 12 identified NBS resistance genes, nine were found to contain a NB-ARC domain. NB-ARCs have been demonstrated to trigger hypersensitive response (HR) leading to localised plant cell death and thus containment of the pathogen (Hammond-Kosack and Jones, 1996; van der Biezen and Jones, 1998). HR occurs in response to pathogen derived molecules (Avr genes) with trigger specificity controlled by LRR domains of the resistance gene (van der Biezen and Jones, 1998). A high frequency of NB-ARC association with Verticillium resistance focal SNPs suggests that HR may play a large role in *V. dahliae* resistance response of strawberry. HR resistance is typically considered to be race specific and also have a lower durability within the field (Lindhout, 2002). Previous studies have highlighted the importance of the HR in roots: *Phytophthora sojae* resistance was partially induced in soybean through the use of lesion mutant lines which triggered root cell death in response to pathogen invasion (Kosslak et al., 1996). This also resulted in a trade-off where lesion mutants exhibited an inability to form symbiotic nodules with nitrogen-fixing bacteria (Kosslak et al., 1996). Further evidence that HR may be an important factor of a resistance response to *V. dahliae* infection can be seen where the effector *PevD1* identified in *V. dahliae* isolated from cotton resulted in HR when infiltrated onto tobacco (Wang et al., 2012) and similarly with the Verticillium effector *Ave1* in tobacco (Fan et al., 2018). Of particular interest was the marker *FaRVd7B3* where the closest resistance gene shows 90% identity to 47% of the resistance gene *muRdr1* controlling gene-for-gene specific resistance to *Diplocarpon rosae* a foliar fungal disease in tetraploid rose (*Rosa multiflora*) (Terefe-Ayana et al., 2011).

### Automated phenotyping as a tool for breeders

The UAV and imaging have allowed the development of a high-throughput phenotyping system to assess the disease resistance status of plants. A substantial labour-saving cost could be achieved through implementation of the phenotyping platform as the manual assessment of 2500 plants five times over the season took a total of 37.5 hr. In contrast switching to a UAV-based phenotyping approach cut the time down to 2.5 hours. There was a strong association between the manual disease scores (AUDPC) and the automated disease scores (NDVI-AUDPC) of *V. dahliae* inoculated plants. Furthermore, the use of automated phenotypic scores resulted in successful identification of resistance markers. In a similar study NDVI was found to be a good measure for Verticillium wilt structural damage in olive (Calderón et al., 2013), which suggests the transferability of this NDVI disease score across different crop hosts. In future work, the semi-automated image analysis will be improved to fully automated canopy segmentation.

### Deploying the identified resistance

Most of the alleles identified in this study are of moderate effect with two out of 25 consistently preforming over the wider germplasm. Studying the Verticillium resistance present within pertinent cultivars related to breeding populations will ensure greater relevance of future resistance markers. In the absence of robust markers associated with the moderate resistance incidences seen here, and in the complete absence of major single gene resistance, we believe that genomic selection may provide a better strategy to breed Verticillium disease resistance into strawberry. Nonetheless, recent advances in strawberry research including recent advances in genome sequencing (unpublished observation) and successful CRIPSR/CAS9 transformation (Wilson et al. 2018), could be used to identify putative resistance genes and allow functional characterisation, respectively. These tools may allow the development of robust functional markers which perfectly tag the causative resistance genes associated with *FaRVd5D* and *FaRVd3D*.

## Conclusions

Marker-assisted breeding and more likely genomic selection will result in a higher probability of developing a successful cultivar containing Verticillium wilt resistance and provides plant breeders with a competitive advantage in comparison to those implementing empirical breeding strategies. Here we report multiple loci of interest for breeders, two of which are associated with resistance across the wider strawberry germplasm. Furthermore, we highlight the potential for a HR resistance mechanism to play a large role in resistance to Verticillium in strawberry. The automated phenotyping platform could provide a valuable tool for breeders and pre-breeding research work.

## Supporting information

## Acknowledgements

The authors acknowledge funding from the Biotechnology and Biological Sciences Research Council (BBSRC) BB/E007074/1 and Innovate UK project 101630. The authors acknowledge the help of Rafael M. Jiménez-Diaz for help with determining the subclade of the *Verticillium dahliae* isolates. The authors acknowledge Eric Van de Weg, Phillip Brain and Xiangming Xu for statistical advice. The authors acknowledge Outfield for drone imaging. The authors also acknowledge Dr Beatrice Denoyes, INRA and Dr Amparo Monfort, CRAG for granting the use of their informative markers in the production of the strawberry consensus linkage map.

## Abbreviations

AUDPC: Area Under the Disease Progression Curve cfu - colony forming units
ExF: ‘Emily’ x ‘Fenella’ mapping population
*FaRVd\*\**: *Fragaria ananassa* Resistance allele for *Verticillium dahliae* ** denotes chromosome (1-7) and subgenome (A-D)
FxC: ‘Flamenco’ x ‘Chandler’ mapping population
GCA: General Combining Ability
HR: Hypersensitive Response
i35k: Istraw35 Affymetrix chip
i90k: Istraw90 Affymetrix chip
NB-ARC: *domain name*
NBS: Nucleotide Binding Site
NDVI: Normalized Difference Vegetation Index
NIR: Near InfraRed
QTL: Quantitative Trait Loci
R: Red
rAUDPC: relative Area Under the Disease Progression Curve
RGB: Red Green Blue
RH%: Percentage Relative Humidity
RxH^a^: ‘Redgauntlet’ x ‘Hapil’ mapping population. Cross one (n=169) - screened with mixed inoculum
RxH^b^: ‘Redgauntlet’ x ‘Hapil’ mapping population. Cross two (n=80) - screened with isolate 12158
SCA: Specific Combined Ability
SNP: Single Nucleotide Polymorphism
SSR: Simple Sequence Repeat
UAV: Unmanned Aerial Vehicle

## Author Contributions

HMC DS, CE and RJH – Conceived and designed experiments.

HMC and CM-M, AGC – performed all pathogenicity tests.

RJV – Analysed SNP data and made linkage map

AJP – Propagated plant material

HMC - Analysed pathogen data & conducted quantitative genetics analysis

BL – Analysed imaging data

DWS – Provided plant material and phenotyping advice

AA – Gene annotations & Bioinformatics support

HMC, BL and RJH wrote the manuscript with contributions from all authors.

## Conflict of interest statement

On behalf of all authors, the corresponding author states that there is no conflict of interest regarding the publication of this work

---

**Supplementary Figure 1** Area under the disease progression curve for ‘Hapil’, ‘Redgauntlet’ and ‘Chandler’ cultivars. Light grey bars represent plants inoculated with Verticillium dahliae isolate 12008 from subclade II-2, white bars 12158 from subclade II-1 and dark grey represents mock inoculated plants

**Supplementary Figure 2** Area under the disease progression curve for each genotype from the ‘Flamenco’ x ‘Chandler’ and ‘Emily’ x ‘Fenella’ populations

**Supplementary Figure 3** Frequency of markers sampled at random which are within 100 kb of a nucleotide binding site (NBS) containing resistance gene. The red vertical line represents the number of markers within 100 kb of an NBS observed in this study.

**Supplementary Figure 4** Frequency of markers sampled at random which are within 100 kb of an NB-ARC containing resistance gene. The red vertical line represents the number of markers within 100 kb of an NB-ARC observed in this study.

**Supplementary Figure 5** Linkage map displaying marker positions (grey) in Mb for 28 linkage groups of octoploid strawberry (1A-7D) marker positions scaled to *F. vesca* genome. Resistance marker locations from the Istraw90 Affymetrix chip (red) and Istraw35 Affymetrix chip validation SNPs (green).

**Supplementary File 1** ‘Flamenco’ x ‘Chandler’ Axiom^®^ IStraw90 marker order based upon *Fragaria vesca* Hawaii 4 genome version 2.0 22.

