## Supplementary figures and images for "Image-based Phenotyping and Disease Screening of Multiple Populations for resistance to *Verticillium dahliae* in cultivated strawberry *Fragaria x ananassa*"

### Supplementary file 1

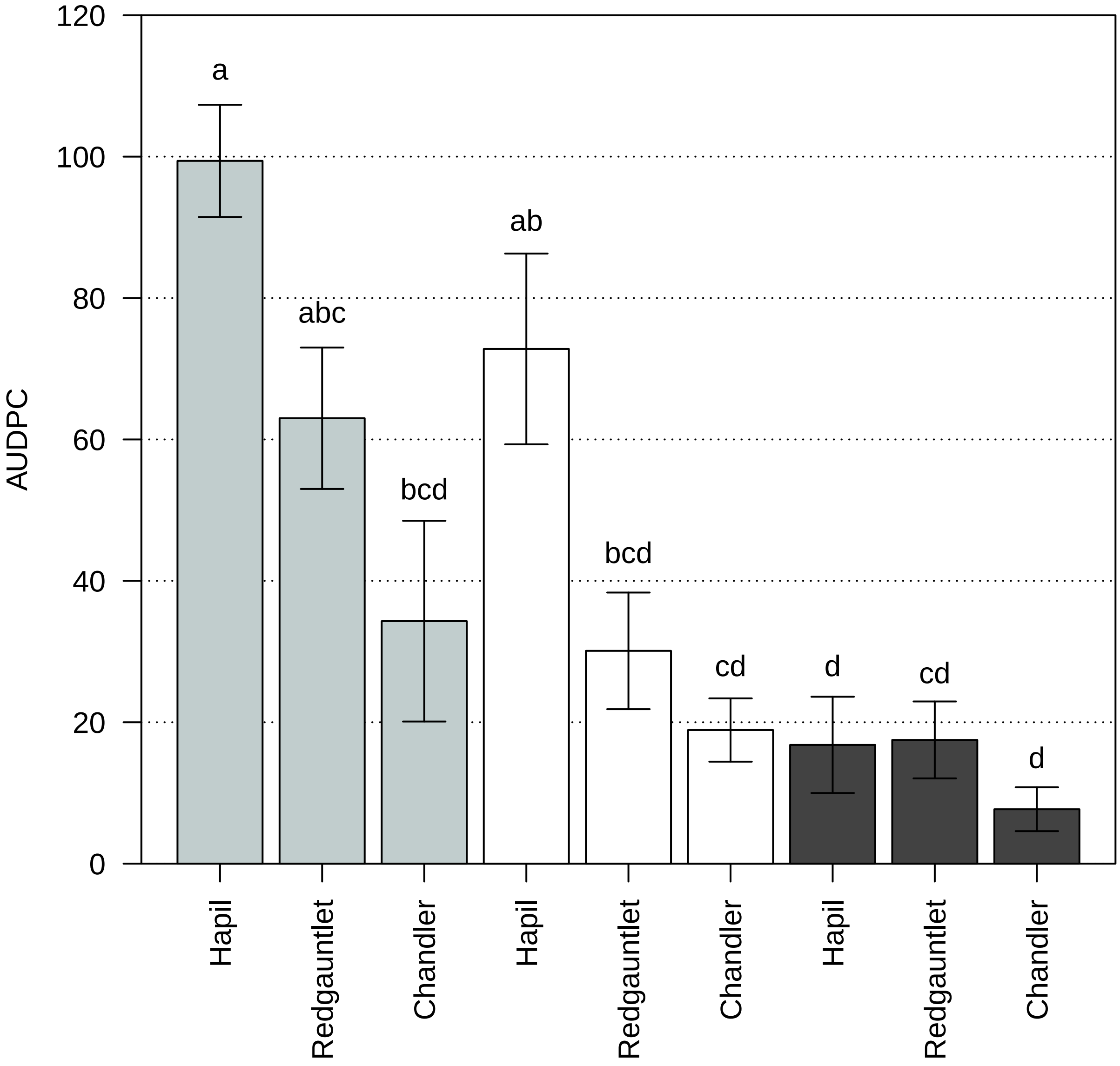

### Supplementary file 2

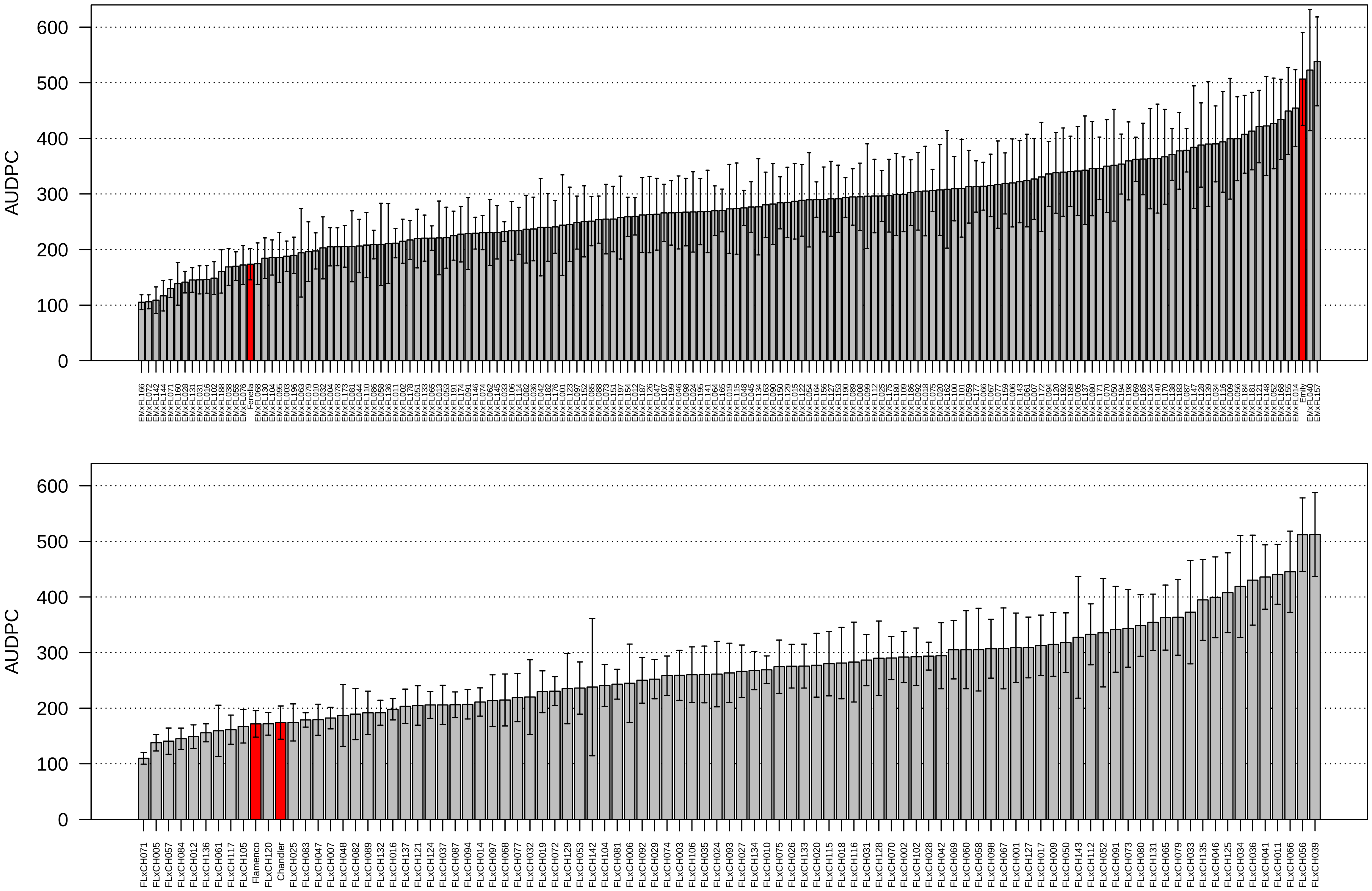

### Supplementary file 3

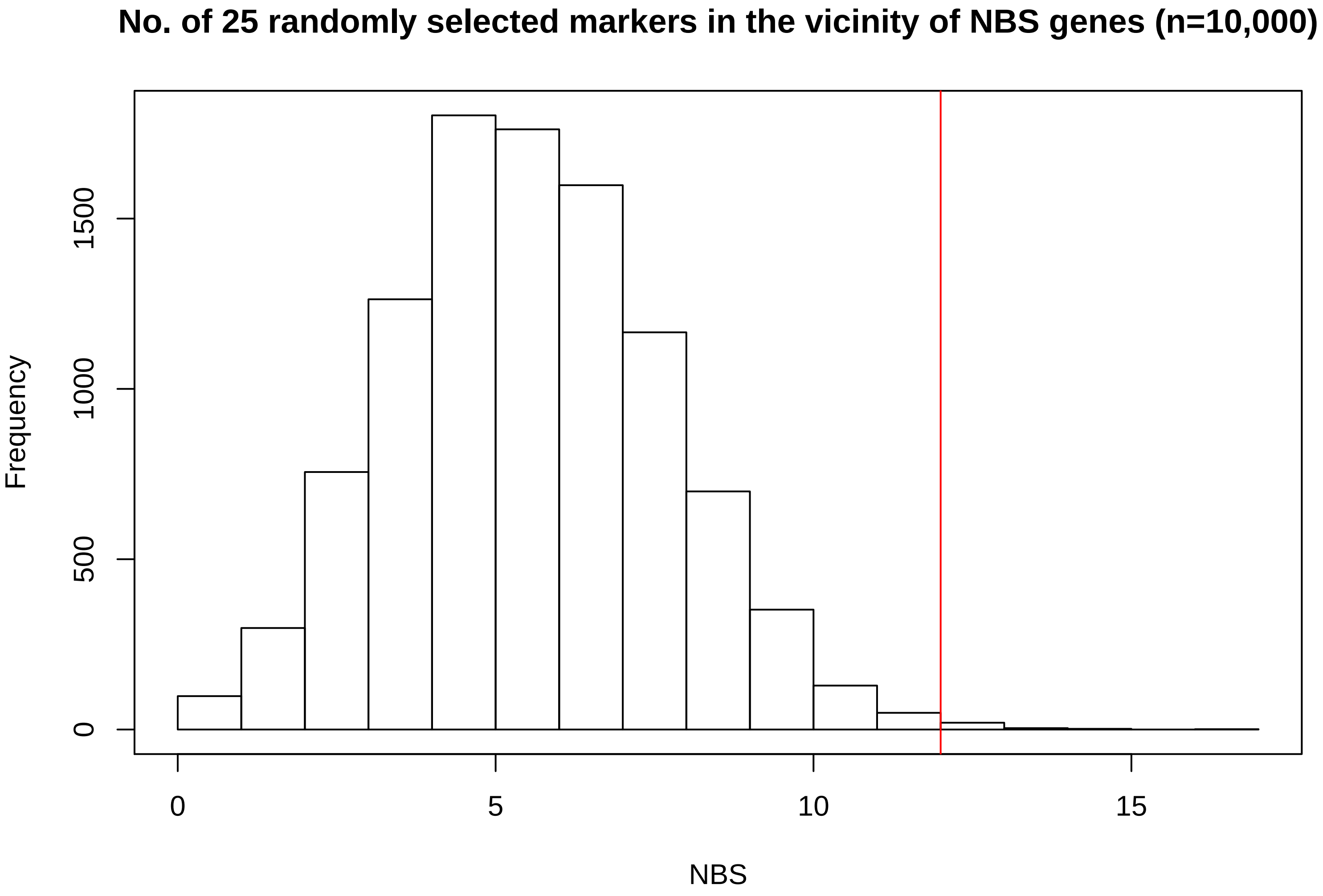

### Supplementary file 4

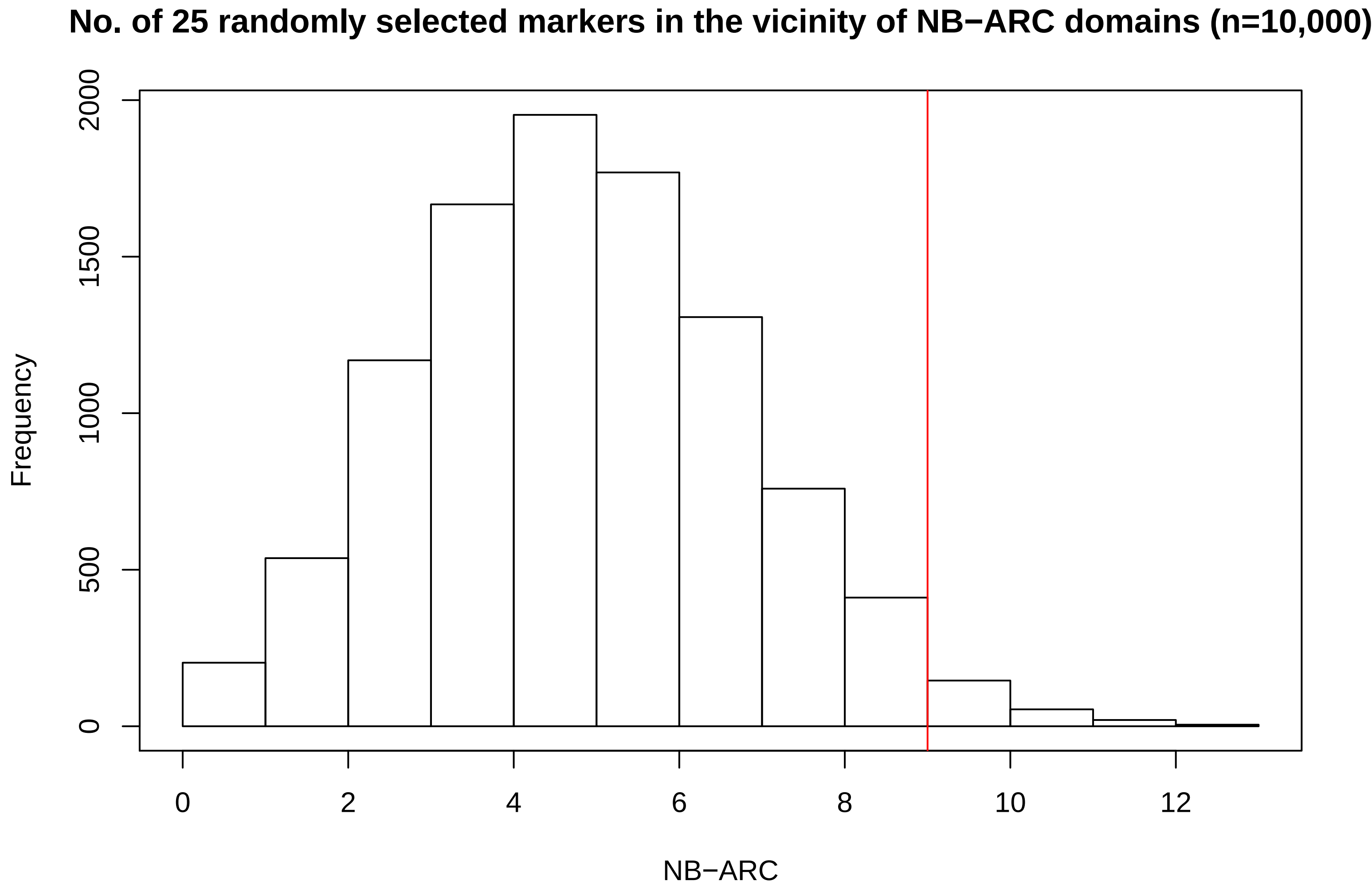

### Supplementary file 5

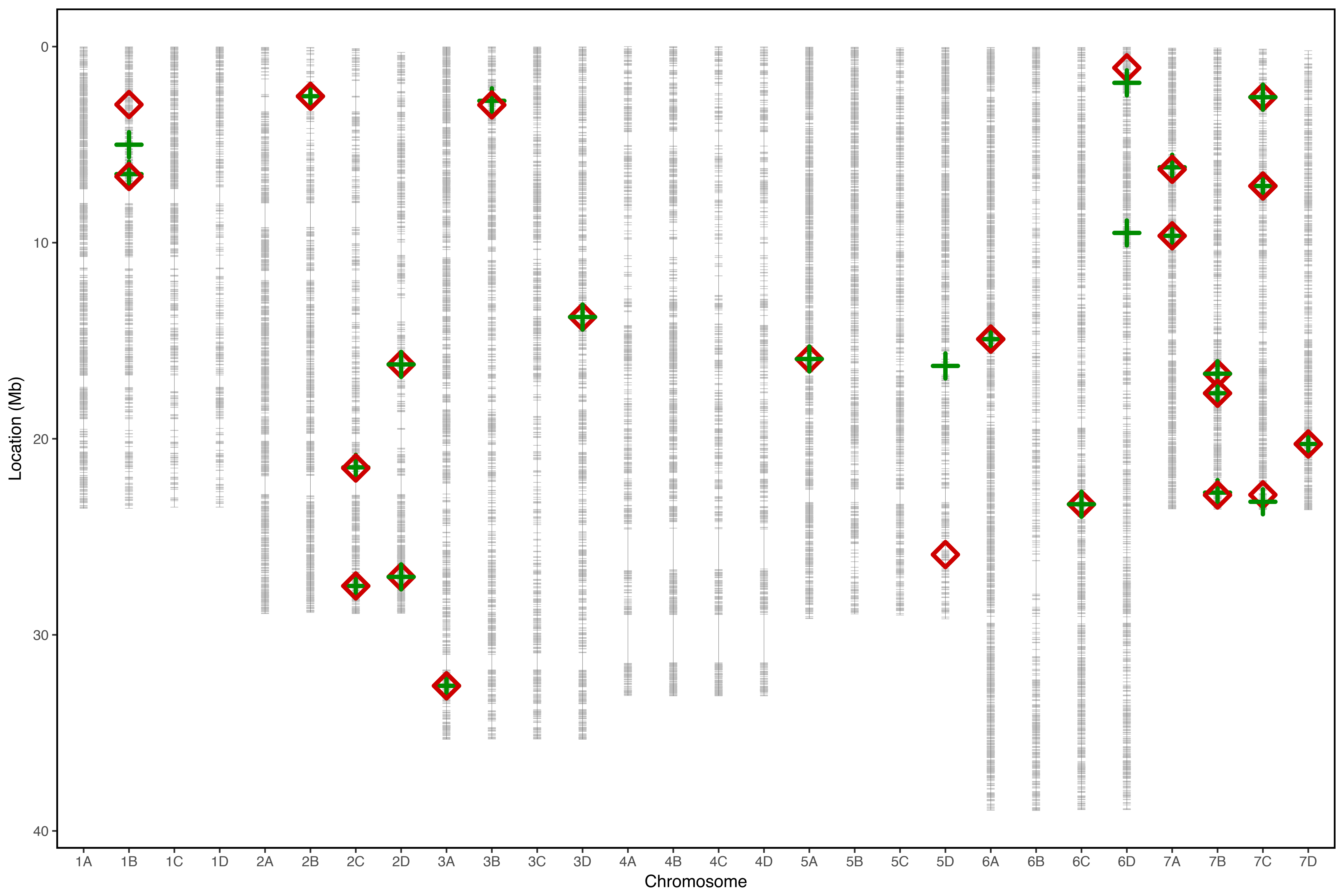
